# Enhanced SNP Genotyping with Symmetric Multinomial Logistic Regression

**DOI:** 10.1101/2024.11.28.625807

**Authors:** Malte B. Nielsen, Poul S. Eriksen, Helle S. Mogensen, Niels Morling, Mikkel M. Andersen

## Abstract

In genotyping, determining Single Nucleotide Polymorphisms (SNPs) is standard practice, but it becomes difficult when analysing small quantities of input DNA, as is often required in forensic applications. Existing SNP genotyping methods, such as the HID SNP Genotyper Plugin (HSG) from Thermo Fisher Scientific, perform well with adequate DNA input levels but often produce erroneously called genotypes when DNA quantities are low. To mitigate these errors, genotype quality can be checked with the HSG. However, enforcing the HSG’s quality checks decreases the call rate by introducing more no-calls, and it does not eliminate all wrong calls. This study presents and validates a Symmetric Multinomial Logistic Regression (SMLR) model designed to enhance genotyping accuracy and call rate with small amounts of DNA. Comprehensive bootstrap and cross-validation analyses across a wide range of DNA quantities demonstrate the robustness and efficiency of the SMLR model in maintaining high call rates without compromising accuracy compared to the HSG. For DNA amounts as low as 31.25 pg, the SMLR method reduced the rate of no-calls by 50.0% relative to the HSG while maintaining the same rate of wrong calls, resulting in a call rate of 96.0%. Similarly, SMLR reduced the rate of wrong calls by 55.6% while maintaining the same call rate, achieving an accuracy of 99.775%. The no-call and wrong-call rates were significantly reduced at 62.5–250 pg DNA. The results highlight the SMLR model’s utility in optimising SNP genotyping at suboptimal DNA concentrations, making it a valuable tool for forensic applications where sample quantity and quality may be decreased. This work reinforces the feasibility of statistical approaches in forensic genotyping and provides a framework for implementing the SMLR method in practical forensic settings. The SMLR model applies for genotyping biallelic data with a signal (e.g. reads, counts, or intensity) for each allele. The model can also improve the allele balance quality check.

## 1. Introduction

In forensic genotyping, the accurate calling of Single Nucleotide Polymorphisms (SNPs) is crucial but becomes a challenge when dealing with low amounts of DNA, which is typical for biological traces. While standard tools, such as the HID SNP Genotyper Plugin (HSG) from Thermo Fisher Scientific (Waltham, MA, USA), excel with sufficient DNA amounts, their performance declines with lower DNA quantities, resulting in increased erroneous genotype calls and reduced call rates.

To reduce the number of wrong calls (WCs), it seems natural to declare a no-call (NC) for genotypes not passing the quality checks (QCs) provided by the HSG. However, for low amounts of input DNA, this approach significantly decreases the call rate and still struggles to filter out WCs.

We introduce a Symmetric Multinomial Logistic Regression (SMLR) model designed to overcome the challenges of low DNA amounts. The SMLR model employs a statistical methodology for refined NC declaration, improving accuracy and call rate, especially in challenging conditions.

We explore the SMLR model’s efficiency and reliability, laying out guidelines for its application in forensic SNP genotyping. We also demonstrate the SMLR model’s potential in quality control, particularly in identifying and managing allelic imbalance.

## 2. Material and Methods

For the analyses, we used *R* version 4.4.2 with the packages: *tidyverse, future*.*apply*, and *xtable* [1–7]. For creating figures, we used *ImageMagick* and the R packages: *ggplot2, ggnewscale, latex2exp*, and *patchwork* [8–12]. The R scripts used for data analyses and figure generation are available on GitHub and Zenodo [13].

### 2.1. The HID SNP Genotyper Plugin

The HSG software uses multiple metrics for SNP genotype determination [14, p. 35]. Along with its genotype calls, it outputs three quality checks:

- A locus-wise coverage check to indicate potential drop-outs (this QC flag was not observed in our data).
- A check of the strand balance where a percentage of positive coverage below 0.3 or above 0.7 results in a QC flag indicating imbalance.
- A check of the allele balance that flags homozygous calls if the ratio of the major allele’s coverage to the total coverage of all four nucleotides falls below 0.95 and heterozygous calls if it falls outside the range of 0.35 to 0.65.

As seen in the leftmost and middle plots of Fig. 1, the HSG’s genotype calls and determination of NCs are not based on these QCs alone, so genotypes are often called despite the presence of QC flags, and NCs are declared even when no QC flag is present [14, p. 35]. Therefore, Enforcing the Quality Checks (EQC) by turning genotypes with QC flags into NCs will increase the number of NCs and thus reduce the call rate.

**Fig. 1:**
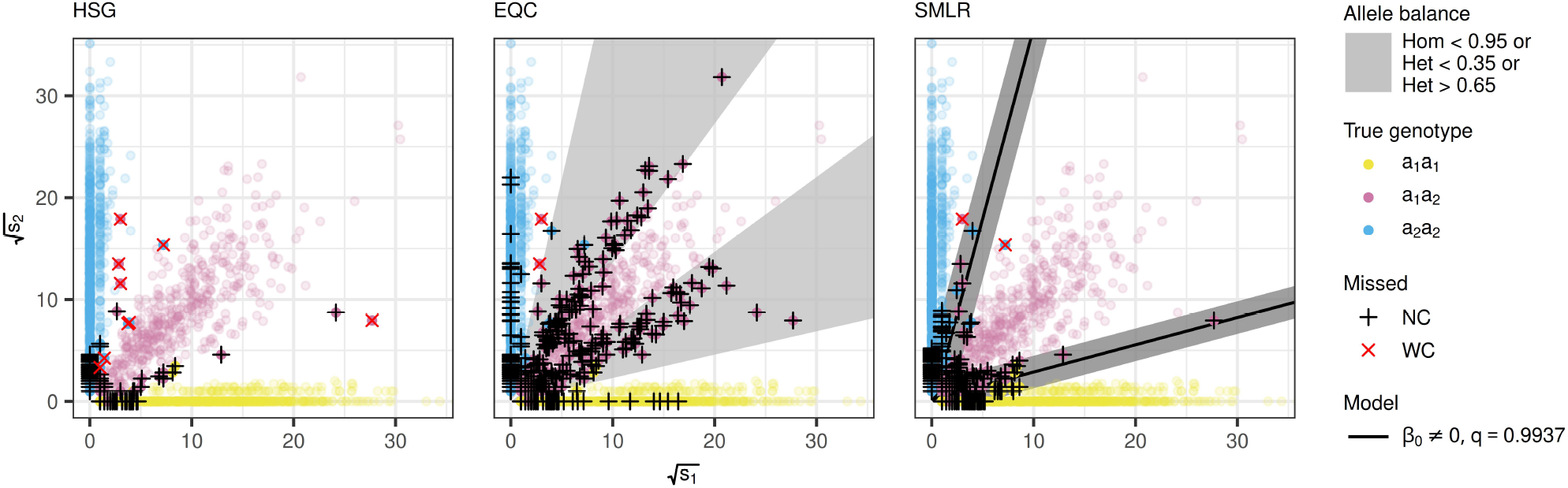
Genotype predictions for the examinations of 31.25 pg DNA. Each plot displays 1,931 SNP observations classified using the genotyping methods: HID SNP Genotyper Plugin (HSG), Enforcing the Quality Checks (EQC), and Symmetric Multinomial Logistic Regression (SMLR). A dot represents a set of read counts (*s*_1_, *s*_2_) for an SNP. It is coloured by the true genotype: red for heterozygous and blue or yellow for homozygous. Failed genotype predictions are indicated by red crosses for wrong calls and black pluses for no-calls. In the EQC plot (middle), the grey areas show where the HSG is guaranteed to flag for allelic imbalance. In the SMLR plot (right), the solid lines show the decision boundaries of the SMLR model with an intercept fitted to square-root transformed allele signals, while the grey area marks the no-call zone when *q* = 0.9937.

### 2.2 The Symmetric Multinomial Logistic Regression (SMLR) Model

#### 2.2.1 Model Formulation

The SMLR model presents a statistical solution to biallelic genotyping challenges by using the allele signals to estimate posterior genotype probabilities. For a biallelic marker with alleles *a*_1_ and *a*_2_ having measured signals *s*_1_ and *s*_2_, we consider the genotype *G* with possible outcomes {*a*_1_*a*_1_, *a*_1_*a*_2_, *a*_2_*a*_2_} and posterior genotype probabilities

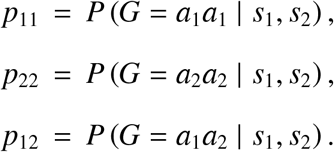

Multinomial logistic regression is apt for modelling the conditional distribution of *G* given *s*_1_ and *s*_2_ with the heterozygous genotype as a baseline category for convenience and standardisation [15, p. 293]. However, it is desirable to model *P*(*G* | *s*_1_, *s*_2_) in a way that is invariant to the labelling of the alleles by introducing a symmetry into the model equations, leading to the SMLR model:

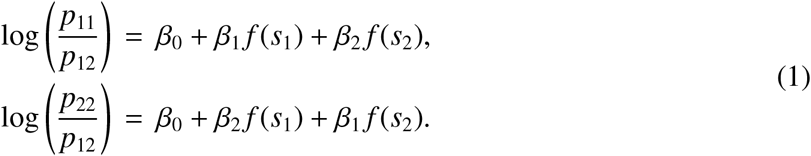

Here, the function *f* is a variance-stabilising transformation of the allele signals, e.g. 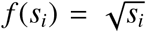.In standard multinomial logistic regression, the second equation would have different β_*i*_-parameters than the first. The introduction of the above symmetry ensures that it does not matter which of the two alleles is labelled *a*_1_ or *a*_2_. The posterior genotype probabilities become

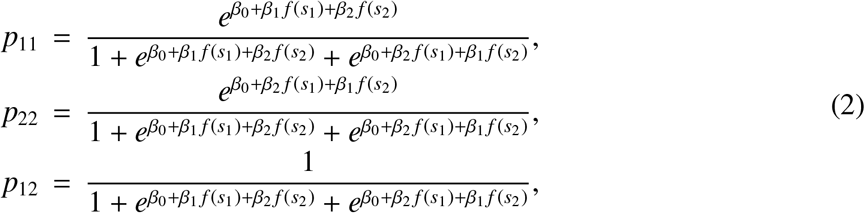

where *β*_1_ is expected to be positive, such that *p*_11_ increases with *s*_1_ and *p*_22_ increases with *s*_2_, and *β*_2_ is expected to be negative, so that *p*_11_ and *p*_22_ decrease with increasing *s*_2_ and *s*_1_, respectively. It is expected that *β*_1_ < |*β*_2_| such that *p*_12_ goes towards 1 as *s*_1_ = *s*_2_ grows large, aligning with the behaviour of heterozygous genotypes at high signal levels.

#### 2.2.2 SNP Genotype Calling

With the SMLR model, the genotype calling for an observation is straightforward: the genotype to be called is that associated with the highest posterior probability estimated from (2). The decision boundaries are the points (*s*_1_, *s*_2_) where at least one of the model equations from (1) equals zero. In other words, where the posterior probability of the heterozygous genotype equals the posterior probability of one of the homozygous genotypes. As shown in the SMLR plot in Fig. 1, these boundaries can be represented graphically in a plot of *f* (*s*_2_) versus *f* (*s*_1_) by the lines

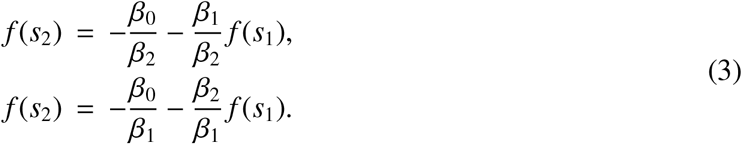

The plot also illustrates how a measure of call confidence is implemented by introducing the user-defined probability threshold, *q*. An observation is then declared an NC if its maximum posterior probability among the potential genotypes falls short of *q*. By setting 1/2 < *q* < 1, observations on the decision boundaries are guaranteed to be declared NCs, yielding unambiguous call decisions even when multiple genotypes have equal posterior probability estimates. The call confidence increases with increasing *q*-values, corresponding to expanding a no-call zone around the decision boundaries within which all posterior genotype probabilities are below *q*.

#### 2.2.3 Estimation

In multinomial logistic regression, maximum likelihood estimates (MLEs) of parameters are well-defined and unique when the data categories overlap and thus are not completely separable [16, 17]. When fitting SMLR models to biallelic data, overlapping categories essentially mean that after applying the variance-stabilising transformation to the allele signals, at least one of the collections of homozygous points cannot be separated from the collection of heterozygous points by any straight line (see SMLR-plot in Fig. 1). For completely separable data, the true genotypes can be perfectly partitioned by the decision boundaries, which implies that multiple versions of the lines from (3) will be equally good in classifying the data. In such cases, the MLEs tend towards infinity unless constrained by a stopping criterion [15, p. 298].

The MLEs are determined by minimising the negative log-likelihood of the SMLR model derived in the Supplementary Material (S1). Using the R function ‘optim’ with default settings facilitates this process. It requires an initial guess for the parameter vector, which can be set to

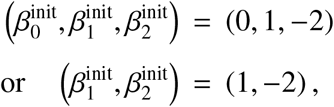

reflecting the expectations of *β*_2_ < 0 < *β*_1_ < |*β*_2_|. These are merely expectations of the resulting parameters, not requirements or constraints on the model equations. Therefore, the initial guess can take values beyond these expectations, such as (0, 0, −1).

### 2.3 Assessing the Minimal Sample Size

Bootstrap analysis was applied to determine a reasonable sample size for fitting the SMLR model. By observing how the variance of parameter estimates decreases with increasing sample size, we identified the optimal number of SNP observations or individuals needed to achieve significant precision gains without excessive data collection [18].

The bootstrap analysis also demonstrates the stability of the SMLR model’s decision boundaries in scenarios of complete separation, which often occurs with smaller sample sizes, and it illustrates under which conditions complete separation is less likely.

### 2.4 Assessing the Effectiveness of the SMLR Model

Cross-validation was used to assess the effectiveness of the SMLR model in increasing the call rate without compromising accuracy and vice versa. In this article, call rate (*CR*) is defined as the percentage of calls that are not NCs, and accuracy (*AC*) as the percentage of correct calls when disregarding NCs:

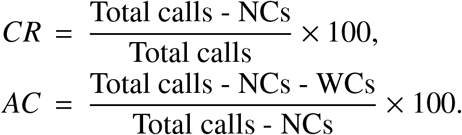

To ensure fair comparisons between the SMLR model and the HSG, it is critical to analyse their performances under equivalent conditions. Therefore, the relative difference in call rates was assessed when the no-call zone of the SMLR model had a width providing the same level of accuracy as the HSG. Since it is not always possible to find a width that gives exactly the same accuracy for the SMLR model and the HSG, the accuracy of the SMLR model was set to the lowest value exceeding the accuracy of the HSG, i.e. its no-call zone was adjusted to the width where the model yielded the same number of WCs as the HSG or less. Conversely, the accuracies were compared when the SMLR model provided at least the same call rate as the HSG, i.e. the same number of NCs or fewer. This way, the comparisons are conservative by favouring the HSG over the SMLR model.

When aligning the SMLR model’s WCs and NCs according to the HSG, the relative differences in call rates and accuracies are proportional to the relative differences in NCs and WCs, respectively:

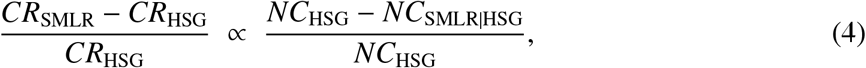

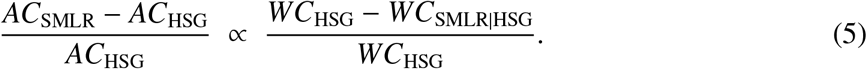

Here, *NC*_HSG_ and *WC*_HSG_ are the no-calls and wrong calls of the HSG, while *NC*_SMLR|HSG_ and *WC*_SMLR|HSG_ are the corresponding counts for the SMLR model when aligned to match the HSG’s WC and NC levels, respectively.

For signals exhibiting variation consistent with a Poisson distribution, commonly observed for integer-valued data, the transformation 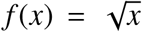 is well established as an effective method for stabilising variance [19]. However, even a theoretically well-justified choice of *f* will not necessarily optimise the performance metrics in (4) and (5). Thus, to explore the effectiveness of various transformations and intercept configurations, six SMLR model formulations were tested: identity, square root, and logarithmic transformations

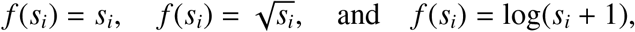

each with and without an intercept (i.e. *β*_0_ ≠ 0 and *β*_0_ = 0).

The models were evaluated through extensive cross-validation, a method that randomly divides the data into disjoint training and test subsets, the former used for model fitting and the latter exclusively to evaluate model performance [20, 21]. Repeating this process with different data splits helps estimate how the model will perform on new datasets. To assess robustness and generalisability, several cross-validations were conducted, considering cases where training and test subsets were drawn from the same DNA dilutions and different dilutions.

This structured approach enables comparing the models in an objective setting and ultimately allows for identifying the optimal model in forensic genetic contexts.

### 2.5 Data

This study analysed results of SNP typing with the Precision ID Ancestry Panel (Thermo Fisher Scientific), which includes 165 autosomal SNPs used to predict the biogeographic origin of humans. The laboratory methods were described by Pereira et al. [22].

The manufacturer recommends using 1ng DNA to increase the success rate with degraded DNA from compromised tissue samples. Others have demonstrated that good results can be obtained with smaller amounts of DNA if it is of good quality and modified experimental conditions are used [23]. Two DNA dilution series were analysed: The first series included six two-fold DNA dilutions from each of five individuals, with 1 ng, 500 pg, 250 pg, 125 pg, 62.5 pg, and 31.25 pg DNA, repeated in four examinations. Due to limited amounts of DNA, the dilution levels of 62.5 pg and 31.25 pg DNA did not include all five individuals. This series included six complete SNP profile measurements for all individuals, with additional partial data. For each individual, these complete SNP profiles were identical and were adopted as the true genotypes for subsequent analyses.

The second series was a single examination of four two-fold DNA dilutions from each of 18 individuals with 50 pg, 25 pg, 12.5 pg, and 6.25 pg DNA. The individuals’ SNP profiles were known from previous analyses.

The project is registered in the University of Copenhagen’s joint record of biobanks and record of research projects containing personal data (514-1056/24-3000) and complies with the rules of the General Data Protection Regulation (Regulation (EU) 2016/679).

#### 2.5.1. Initial SNP Calling

The primary sequencing analysis was performed with Torrent Suite Software v4.6 (Thermo Fisher Scientific). BAM files were generated using the HSG (v4.3.1). No noise filter was used.

Some SNPs are known to have rare variants. Therefore, it was investigated whether rare alleles, different from the two expected alleles, were present. The HSG does this through its QC of the allele balance [14, p. 35]. One marker, rs7722456, exhibited anomalous adenine reads and was excluded from the main analyses, along with the markers rs459920 and rs7251928, as recommended by Pereira et al. [22], leaving data from 162 SNPs for analysis. However, rs7722456 was retained as example data to illustrate how the SMLR model can enhance the detection rate of imbalance when used for QC of the allele balance.

SNP data with zero reads for both alleles were removed from the dataset, as they provided no useful information and consistently resulted in NCs for the HSG.

## 3. Results

For low amounts of DNA, the HSG-plot and EQC-plot in Fig. 1 reveal that using the QC-flags to improve the accuracy is inefficient: converting flagged observations to NCs has a significant cost to the call rate and does not even eliminate all WCs. Hence, any meaningful comparison of the SMLR model should be made with the HSG alone. The ensuing results affirm the SMLR model’s utility in forensic genotyping, demonstrating its capacity to improve call rate and accuracy across various testing scenarios.

### 3.1 Pre-Analysis

Table 1 demonstrates how a fit of the SMLR model with an intercept and a square-root transformation reduces NCs across all examined DNA amounts when compared to the HSG using (4) and similarly reduces WCs across all DNA amounts when compared using (5). Thus, for each examined DNA amount, this specific fit of the SMLR model yields fewer NCs and WCs when the *q*-threshold is set to a value between those displayed in Table 1, implying simultaneous improvements in call rate and accuracy. Since the metrics in (4) and (5) favour the HSG, and the cross-validations presented later show that the fit used in Table 1 is not necessarily the best performing, the improvements displayed in the table are conservative.

**Table 1:** The SMLR model’s genotyping improvements for all examined DNA quantities.

|  |  | First series: |  |  |  |  |  | Second series: |  |  |  |
| --- | --- | --- | --- | --- | --- | --- | --- | --- | --- | --- | --- |
|  |  | 5 individuals, 4 examinations |  |  |  |  |  | 18 individuals, 1 examination |  |  |  |
| DNA quantity (pg) |  | 1,000 | 500 | 250 | 125 | 62.5 | 31.25 | 50 | 25 | 12.5 | 6.25 |
| Number of SNPs |  | 3,240 | 3,240 | 3,240 | 3,238 | 2,585 | 1,931 | 2,831 | 2,821 | 2,575 | 2,347 |
| HSG* | No-calls | 1 | 3 | 28 | 48 | 83 | 156 | 62 | 142 | 213 | 276 |
|  | Wrong calls | 0 | 0 | 1 | 1 | 2 | 9 | 8 | 59 | 184 | 272 |
| $q$ aligning WCs | | 0 | 0 | 0 | 0.5473 | 0.5473 | 0.7394 | 0.8115 | 0.8449 | 0.8302 | 0.8541 |
| SMLR<sup>†</sup> | Call rate (%) | 100 | 100 | 100 | 99.8 | 99.7 | 96.0 | 98.4 | 95.7 | 95.1 | 92.2 |
|  | No-calls<sup>‡</sup> | 0 | 0 | 0 | 5 | 7 | 78 | 46 | 121 | 125 | 184 |
|  | NC-reduction (%)<sup>§</sup> | 100 | 100 | 100 | 89.6 | 91.6 | 50.0 | 25.8 | 14.8 | 41.3 | 33.3 |
| $q$ aligning NCs | | 0.8890 | 0.8808 | 0.8588 | 0.8592 | 0.8514 | 0.8414 | 0.8488 | 0.8592 | 0.8725 | 0.8841 |
| SMLR<sup>‡</sup> | Accuracy (%) | 100 | 100 | 100 | 100 | 100 | 99.775 | 99.747 | 97.878 | 92.760 | 88.048 |
|  | Wrong calls | 0 | 0 | 0 | 0 | 0 | 4 | 7 | 57 | 171 | 248 |
|  | WC-reduction (%)<sup>§</sup> | 0 | 0 | 100 | 100 | 100 | 55.6 | 12.5 | 3.4 | 7.1 | 8.8 |

\* Number of no-calls (NCs) and wrong calls (WCs) made by the HID SNP Genotyper Plugin (HSG).
<sup>†</sup> The SMLR model's performance when $q$ is set to align WCs according to (4).
<sup>‡</sup> The SMLR model's performance when $q$ is set to align NCs according to (5).
<sup>§</sup> Results are conservative and based on the SMLR model with an intercept fitted to square-root transformed allele signals from examinations of 25 pg and 50 pg DNA.

#### Relevant Dilutions

Fig. 2 shows how the distribution of 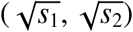 becomes more spread out and how the HSG makes more NCs and WCs as the DNA amount decreases. It further illustrates that most WCs at low DNA amounts are heterozygous genotypes misidentified as homozygous due to a reduced signal for one of the alleles. For the SMLR model, this pattern necessitates higher *q*-threshold settings for accurate genotyping, resulting in lower call rates. Consequently, genotyping with the Precision ID Ancestry Panel requires more than 25 pg DNA to maintain acceptable call rates.

**Fig. 2:**
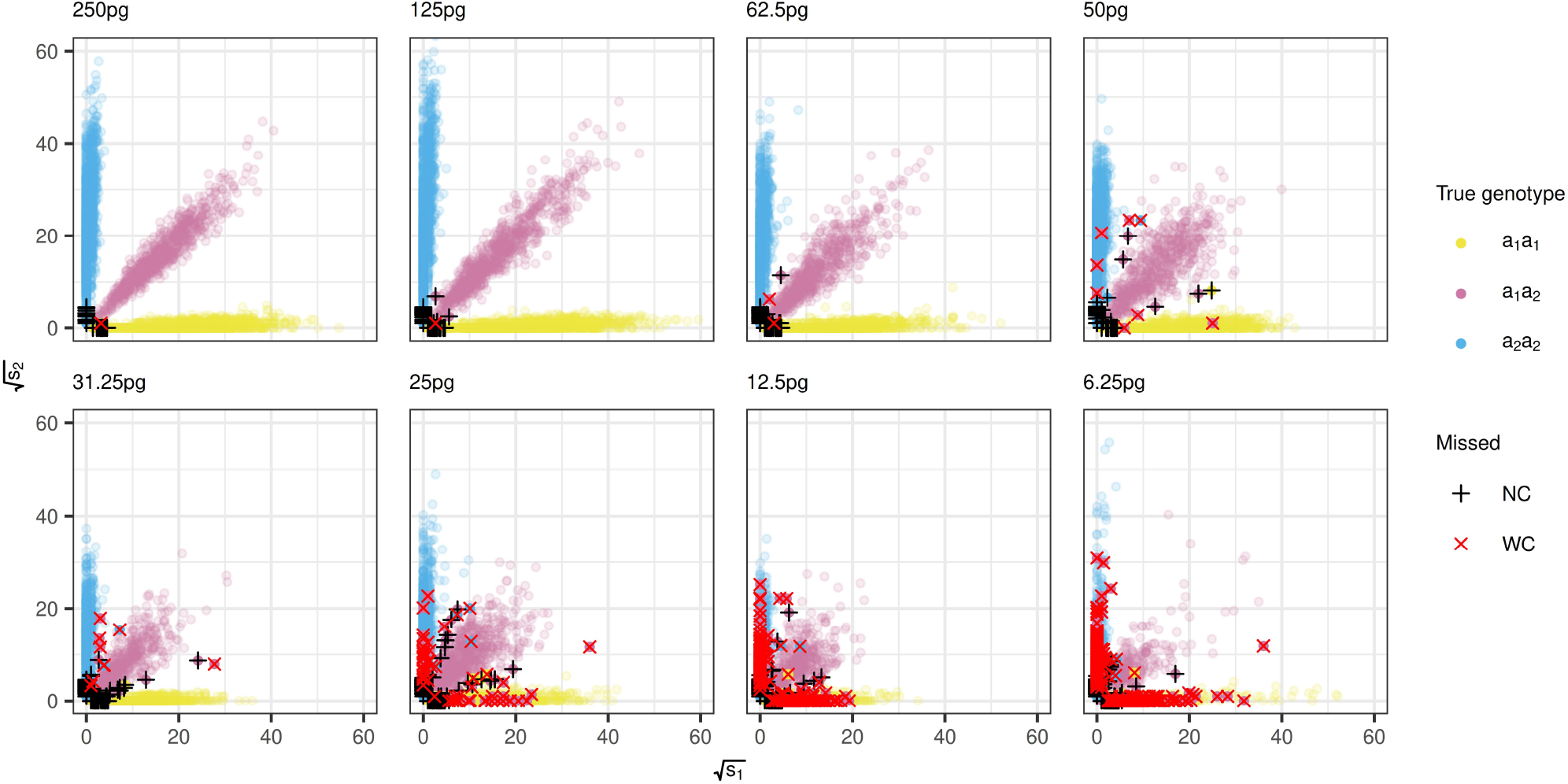
Distribution of square-root transformed allele signals. A dot represents a set of read counts (*s*_1_, *s*_2_) for an SNP and is coloured according to the true genotype: red for heterozygous and blue or yellow for homozygous genotypes. The displayed DNA quantities indicate where the accuracy of the HID SNP Genotyper Plugin falls below 100%, with its wrong calls marked by red crosses and no-calls by black pluses.

Table 1 highlights performance variations across different DNA input levels. At 500 pg DNA and above, the HSG performs adequately, rendering the SMLR model redundant. At 250 pg DNA, the HSG produces a few WCs and introduces significantly more NCs with each step down the dilution series. Fig. 2 and Table 1 show that the HSG makes significantly more WCs at 50 pg DNA and below.

Based on these observations, the following analyses focus on testing genotyping performance within the two ranges of DNA quantities: 31.25–50 pg and 62.5–250 pg. The former is where the call rate of the HSG becomes critically low and it more frequently makes WCs. However, to assess robustness and generalisability, the SMLR models will also be fitted using data from exterior DNA quantities.

#### Parameter Variance, Sample Size, and Separation

Bootstrap analyses were performed for the six SMLR variants mentioned in the Material and Methods section. They all showed similar results to those depicted in Supplementary Fig. S1. Here, we see a rapid decline in the parameters’ variances until a sample size of around nine individuals or roughly 1,470 SNPs. This sample size provides a good balance between the proportions of data used for fitting and testing in the cross-validation, especially in the least populated examination (31.25 pg DNA), where the use of nine individuals for fitting corresponds to 75% of the data.

Supplementary Fig. S1 indicates that the parameter estimates and decision boundaries depend on the DNA amount, with the effect becoming more noticeable at 12.5 pg and below. It also shows that complete separation can be avoided by including examinations of very low DNA amounts (e.g. 25 pg) in the fitting data, and that this does not substantially alter the decision boundaries.

### 3.2 Validation of the SMLR Model

#### Cross-Validation Insights

The cross-validation experiments in Supplementary Fig. S2 demonstrate that the strong results from Table 1 were not coincidental. The figure shows that when either the square root or logarithmic transformation is applied to the allele signals, the SMLR model generally outperforms the HSG with substantial reductions in NCs and WCs, improving both call rate and accuracy.

The subplots in the figure’s first and fourth columns show nearly horizontal performance lines, indicating that fitting to combined data from the examinations of 25 pg and 50 pg DNA gives particularly stable SMLR models for both the low (31.25–50 pg) and high (62.5–250 pg) DNA amounts.

Therefore, the subsequent analyses are based on these fits to let the results inherit this stability.

Supplementary Fig. S2 intends to compare each SMLR variant to the HSG and not to make comparisons between the variants themselves, as a variant that excels when aligned to the HSG is not necessarily the best performing for other balances between the call rate and accuracy.

Fig. 3 compares the SMLR variants fitted to data subsets from the examinations of 25 pg and 50 pg DNA: the call rates from the 1,000 cross-validation iterations were rounded to one decimal place, and the median accuracy for each variant was determined at each rounded call rate. This approach allows for a more precise assessment of how the accuracy of each SMLR variant changes with the call rate.

**Fig. 3:**
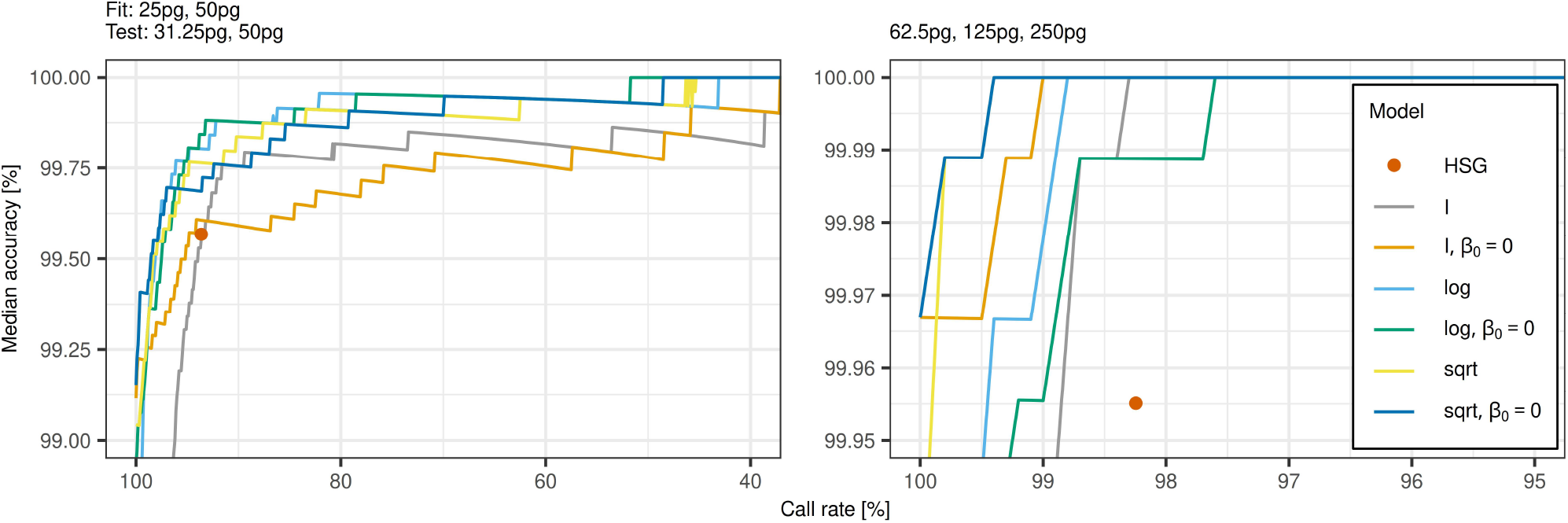
Median accuracy versus call rate from aggregated results of cross-validations. Each line represents the median accuracies calculated from binned call rate data of 1,000 cross-validation iterations, where the six SMLR models were fitted to and tested on data from the examinations of the DNA quantities indicated above the plots. The red dot in each plot shows the median accuracy and call rate for the HID SNP Genotyper Plugin. Note that the first axes have been reversed to align with Fig. 4, where increasing *q* leads to decreasing call rates.

The red dot in each plot of Fig. 3 represents the median accuracy and call rate of the HSG. The lines for the models with 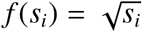 and *f* (*s*_*i*_) = log(*s*_*i*_ + 1) are generally seen to move well to the left of and above these dots. Thus, these four SMLR models convincingly outperform the HSG in median accuracy and call rate. For the low DNA amounts, they increase the call rate to more than 95% while still achieving higher accuracies than the HSG. At the highest call rates, the square-root model without an intercept (dark blue line) achieves slightly higher median accuracies than the other variants.

To avoid overestimating the performance of the SMLR method framework, the remaining analyses were based on the square-root model with an intercept, as this is a more conservative variant given that better SMLR models exist.

#### Call Rate and Accuracy Dependencies

The accuracy of the SMLR model depends on the call rate through the width of the no-call zone, which is controlled by the value of the probability threshold *q*. This dependency on *q* is depicted in Fig. 4, where the previously selected SMLR model is applied to the range of low DNA quantities. The left plot shows that the SMLR model surpasses the HSG’s median accuracy of 99.567% at a probability threshold of *q* = 0.74. At this *q*-value, the right plot shows that the SMLR model has a median call rate of 96.6%, i.e. an increase of 3.0 percentage points compared to the HSG.

**Fig. 4:**
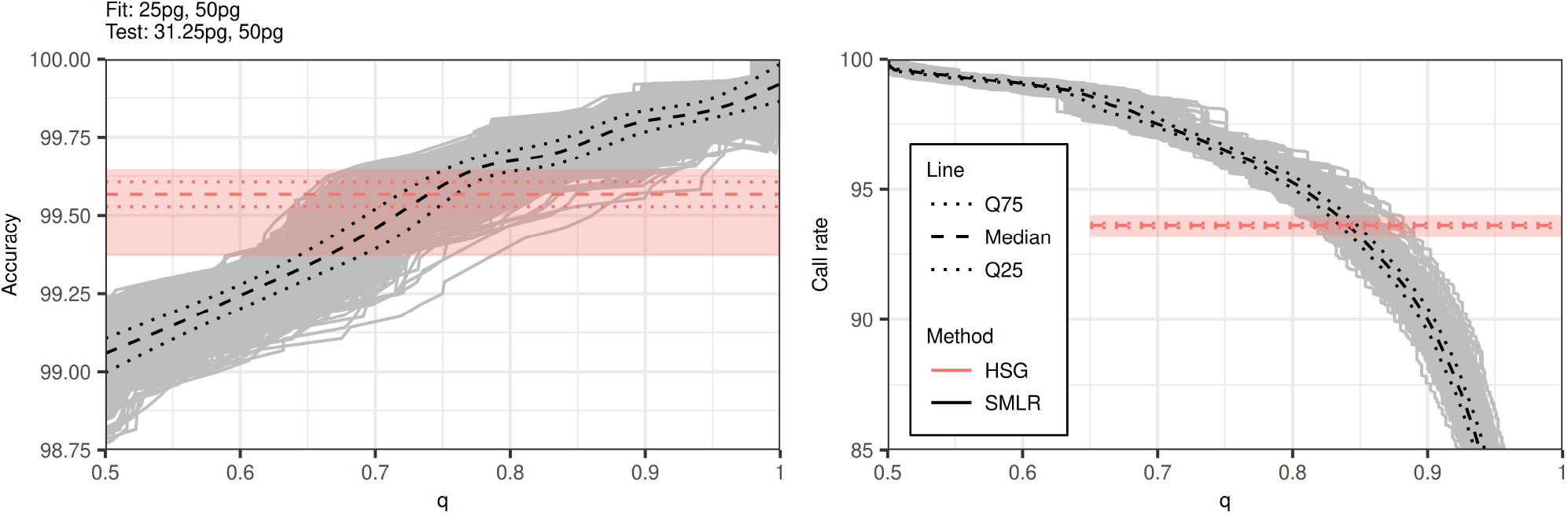
Performance of the SMLR model across probability thresholds, *q*. The grey lines depict the model’s performance in each of 1,000 cross-validation iterations. The model was fitted with an intercept to square-root transformed allele signals and tested on data from the examinations of the DNA quantities indicated above the left plot. The shaded horizontal bars represent the accuracy (left) and call rate (right) ranges for the HID SNP Genotyper Plugin. Dotted lines mark the 25th and 75th percentiles among the 1,000 cross-validations, while dashed lines indicate medians.

The model’s median call rate remains higher than that of the HSG until *q* = 0.84, where it achieves a median accuracy of 99.727%. The left plot (Fig. 4) shows that this is above the maximum accuracy observed for the HSG (99.647%) during the 1,000 cross-validation iterations.

When applied to DNA quantities of 500 pg and above, Table 1 shows that the SMLR model achieves 100% accuracy and call rate. This demonstrates its capability to maintain high performance across varying DNA inputs.

#### SMLR for Quality Checks

The locus rs7722456 had many reads of an unexpected nucleotide (adenine) for all examinations in the first dilution series. The HSG’s quality control flagged only 37 of the 108 unusual observations with its QC flag for allelic imbalance. The limited efficiency in identifying allele balance issues was consistent for all DNA quantities examined in this dilution series.

In Supplementary Fig. S3, the major allele signals for rs7722456 are plotted against the adenine signals, along with the decision boundaries of the previously selected SMLR model. Using the probability threshold *q* = 0.99, 106 points were in the no-call zone, indicating an allelic imbalance. The choice of *q* = 0.99 is a mere example illustrating how the SMLR model can also be used for QC and why this statistical approach is more effective than applying a static threshold to the coverage ratio [14, p. 35].

## 4. Discussion

This study aimed to develop an effective method for SNP calling in forensic genetics. The developed SMLR model is simple, using only two or three parameters to compute three posterior genotype probabilities from two input signals. It is possible to use more complex variance-stabilising transformations than those investigated in this study. Still, the aim was to demonstrate the capabilities of the model while retaining as much of its inherent simplicity as possible.

Measuring the performance of the SMLR variants is not straightforward when NCs are introduced and should be judged neutrally, i.e. from the perspective that they are meant to remove WCs, whereby NCs should not be seen as incorrect genotype predictions. This is further complicated as performance changes with the amount of input DNA. However, the SMLR model using a squareroot transformation emerged as the best all-in-one model, offering a good balance between NCs and WCs for DNA quantities as low as 31.25 pg, particularly when modelled without an intercept.

The SMLR method was compared with the commonly used HSG, which performs well with high DNA quantities but leaves room for improvement at lower DNA amounts. The SMLR models using a square-root transformation generally perform better than the HSG on the examined DNA quantities. Consistently outperforming a genotyping method with high accuracy and call rate, such as the HSG, is a significant achievement that demonstrates the applicability of the SMLR method for biallelic genotyping, particularly in forensic genetics settings.

While the improvements in call rate and accuracy were more modest for the high DNA amounts recommended by kit manufacturers, we demonstrated that the SMLR framework substantially increased the detection rate of allelic imbalance independently of the examined DNA quantity (Supplementary Fig. S3). As a result, any laboratory can benefit from the SMLR framework, regardless of the DNA quantities they typically work with.

Since data characteristics may vary when using SNP platforms and procedures different from those in this study, the SMLR model’s parameters may need adjustment and validation before being implemented with new genotyping methods. It should be noted that the effect of changes in the probability threshold, *q*, on the width of the no-call zone depends on the parameter estimates, particularly on the choice of variance-stabilising transformation. A tailored dilution series can help define the optimal setup for each lab environment, especially in determining the ideal value for the probability threshold *q*.

The SMLR method can be adapted with minor modifications to analyse data from other biallelic genetic systems, e.g. insertion-deletions. The SMLR principle can also classify non-genetic data with similar structures.

The bootstrap analysis showed that using approximately 1,470 SNP observations markedly decreases the parameter estimates’ variances when fitting the SMLR model (Supplementary Fig. S1). For the Precision ID Ancestry Panel, this corresponds to approximately nine complete SNP profiles, with the option of repeating examinations of the same individual, e.g. by measuring the SNP profiles of three individuals, each repeated in three examinations. However, if feasible, we recommend using larger sample sizes to enhance precision further.

For robust fitting, it is important to have a reasonable overlap between the genotype clusters, preferably by measuring sufficient data in the 31.25–50 pg DNA range. If this is not feasible, including a few examinations with as little as 25 pg DNA can effectively achieve overlap. However, the bootstrap analysis showed that adding data from examinations of extremely low DNA quantities, like 12.5 pg, may negatively distort the model’s decision boundaries.

In the Estimation subsection, it was suggested that the initial value of the parameter vector be set to (1, −2) or to (0, 1, −2) if an intercept is included. Depending on the scale of the signals and the choice of variance-stabilising transformation, other initial values may be more suitable. For example, we encountered issues with the initialisation of ‘optim’ when using the suggested initial values for the models with an identity transformation. However, these issues were resolved by setting 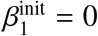 and 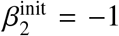.Initialisation problems can occur due to numerical overflow, e.g. if the value of the likelihood function at the chosen initial value exceeds the limit the computer can represent. Such issues are of general concern for optimisation tasks and are not specific to fitting the SMLR model.

Maximum likelihood estimation focuses on optimally placing the decision boundaries based on the observed data but only to classify genotypes. The MLEs can be said to optimise accuracy when the probability threshold, *q*, is set to zero, but when *q* > 1/2, the classification process is changed in a non-trivial way. While the cross-validations demonstrated that the MLEs produce effective SMLR models, it may still be possible to adjust the position of the no-call zone to further improve accuracies and call rates. However, defining an ‘optimal’ position for the no-call zone is not straightforward, as it depends on its width. Nevertheless, research into estimation approaches that incorporate NCs or aim to optimise the position of the no-call zone rather than the decision boundaries could be valuable for forensic genotyping.

Genetic software, such as GenoGeographer [24–26], typically assumes that genotype calls are correct (i.e. without uncertainty) once they pass quality filters. However, genotyping inherently carries some uncertainty, particularly in forensic genetics, where samples of low quality and quantity are common. A statistical approach, such as the SMLR model, provides estimates of posterior genotype probabilities, which can help mitigate this uncertainty. Incorporating these probabilities into tools like GenoGeographer would allow them to account for genotyping errors rather than relying solely on hard calls, thereby improving the accuracy of analyses, particularly when working with DNA samples of low quality and quantity.

## 5. Conclusion

The SMLR model improved the SNP calling efficiency, particularly with suboptimal DNA amounts, as is often the case in forensic genetic examinations of stain material in criminal investigations. At 31 pg DNA, the NC rate was reduced by approximately 50% compared to the HSG, while maintaining the same WC rate as the HSG. The WC rate was reduced by over 50% while maintaining the same NC rate. At 62–250 pg DNA, the no-call rate was dramatically reduced. Using SMLR for quality checks of the allele balance substantially improved imbalance detection regardless of the DNA input level.

## 7. Acknowledgments

We thank Martin Jensen, Sidsel Raaby, and Hai Trieu Nguyen for laboratory assistance, and Martin Tuwel Jensen for technical assistance.

