## Supplementary Material for "Enhanced SNP Genotyping with Symmetric Multinomial Logistic Regression"

*The Negative Log-Likelihood Function*

Let $\mathcal{I}_{11}$ and $\mathcal{I}_{22}$ be index sets for the observations with homozygous genotypes in allele $a_1$ and $a_2$, respectively, and let $\mathcal{I}_{12}$ be the index set for the observations with heterozygous genotypes. Let $m \in \mathcal{I} = \mathcal{I}_{11} \cup \mathcal{I}_{22} \cup \mathcal{I}_{12}$ be a general index and define $t_m = (1, f(s_1^m), f(s_2^m))$, where $s_1^m$ and $s_2^m$ denote the allele signals for $a_1$ and $a_2$ of the observation with index $m$. In accordance with the notation used in (2), let $p_{11}^i$, $p_{22}^j$, and $p_{12}^k$ be the posterior genotype probabilities of the observations with indices $i \in \mathcal{I}_{11}$, $j \in \mathcal{I}_{22}$, and $k \in \mathcal{I}_{12}$, respectively. To ease the notation further, define $\beta = (\beta_0, \beta_1, \beta_2)$ and $\tilde{\beta} = (\beta_0, \beta_2, \beta_1)$. The likelihood for $\beta$ is

$$\mathcal{L}(\beta) = \prod_{i \in \mathcal{I}_{11}} p_{11}^i \prod_{j \in \mathcal{I}_{22}} p_{22}^j \prod_{k \in \mathcal{I}_{12}} p_{12}^k,$$

and the log-likelihood can be written as

$$\begin{aligned}
\ell(\beta) &= \sum_{i \in \mathcal{I}_{11}} \log\left(p_{11}^i\right) + \sum_{j \in \mathcal{I}_{22}} \log\left(p_{22}^j\right) + \sum_{k \in \mathcal{I}_{12}} \log\left(p_{12}^k\right) \\
&= \sum_{i \in \mathcal{I}_{11}} \log\left(\frac{\exp(\beta \cdot t_i)}{1 + \exp(\beta \cdot t_i) + \exp\left(\tilde{\beta} \cdot t_i\right)}\right) \\
&\quad + \sum_{j \in \mathcal{I}_{22}} \log\left(\frac{\exp\left(\tilde{\beta} \cdot t_j\right)}{1 + \exp\left(\beta \cdot t_j\right) + \exp\left(\tilde{\beta} \cdot t_j\right)}\right) \\
&\quad + \sum_{k \in \mathcal{I}_{12}} \log\left(\frac{1}{1 + \exp(\beta \cdot t_k) + \exp\left(\tilde{\beta} \cdot t_k\right)}\right) \\
&= \sum_{i \in \mathcal{I}_{11}} \left[\beta \cdot t_i - \log\left(1 + \exp(\beta \cdot t_i) + \exp\left(\tilde{\beta} \cdot t_i\right)\right)\right] \\
&\quad + \sum_{j \in \mathcal{I}_{22}} \left[\tilde{\beta} \cdot t_j - \log\left(1 + \exp\left(\beta \cdot t_j\right) + \exp\left(\tilde{\beta} \cdot t_j\right)\right)\right] \\
&\quad - \sum_{k \in \mathcal{I}_{12}} \log\left(1 + \exp(\beta \cdot t_k) + \exp\left(\tilde{\beta} \cdot t_k\right)\right).
\end{aligned}$$

The sums over the logarithms can be collected under the same summation sign to obtain

$$\ell(\beta) = \sum_{i \in \mathcal{I}_{11}} \beta \cdot t_i + \sum_{j \in \mathcal{I}_{22}} \tilde{\beta} \cdot t_j - \sum_{m \in \mathcal{I}} \log\left(1 + \exp(\beta \cdot t_m) + \exp\left(\tilde{\beta} \cdot t_m\right)\right).$$

Multiplying both sides by $-1$ and writing out the dot product between the parameters and the signals, the negative log-likelihood becomes

$$
\begin{aligned}
-\ell(\beta) = &\sum_{m\in\mathcal{I}} \log\left(1 + e^{\beta\cdot t_m} + e^{\tilde{\beta}\cdot t_m}\right) - \sum_{i\in\mathcal{I}_{11}} \beta\cdot t_i - \sum_{j\in\mathcal{I}_{22}} \tilde{\beta}\cdot t_j \\
= &\sum_{m\in\mathcal{I}} \log\left(1 + e^{\beta_0+\beta_1 f\left(s_1^m\right)+\beta_2 f\left(s_2^m\right)} + e^{\beta_0\ +\beta_2 f\left(s_1^m\right)+\beta_1 f\left(s_2^m\right)}\right) \\
&- \sum_{i\in\mathcal{I}_{11}} \beta_0 + \beta_1 f\left(s_1^i\right) + \beta_2 f\left(s_2^i\right) \\
&- \sum_{j\in\mathcal{I}_{22}} \beta_0 + \beta_2 f\left(s_1^j\right) + \beta_1 f\left(s_2^j\right) \\
= &\sum_{m\in\mathcal{I}} \log\left(1 + e^{\beta_0+\beta_1 f\left(s_1^m\right)+\beta_2 f\left(s_2^m\right)} + e^{\beta_0\ +\beta_2 f\left(s_1^m\right)+\beta_1 f\left(s_2^m\right)}\right) \\
&- \beta_0|\mathcal{I}_{11} \cup \mathcal{I}_{22}| \\
&- \beta_1\left(\sum_{i\in\mathcal{I}_{11}} f\left(s_1^i\right) + \sum_{j\in\mathcal{I}_{22}} f\left(s_2^j\right)\right) \\
&- \beta_2\left(\sum_{i\in\mathcal{I}_{11}} f\left(s_2^i\right) + \sum_{j\in\mathcal{I}_{22}} f\left(s_1^j\right)\right),
\end{aligned}
\tag{S1}
$$

where $|\mathcal{I}_{11} \cup \mathcal{I}_{22}|$ is the number of homozygous observations. Note how $\beta_1$ is multiplied by a term that ideally sums the major allele signals of the homozygous genotypes, while $\beta_2$ is multiplied by a term that ideally sums the minor allele signals, which are expected to be close to zero.

*Technical Details on the Cross-Validation*

Each of the six columns of plots in Supplementary Fig. S2 represents its own cross-validation. As indicated at the top of the figure, each of the cross-validations used different collections of DNA amounts to fit and test the models. In each cross-validation, the data was split randomly 1,000 times into disjoint pairs of fitting and testing subsets on which all of the models were validated. The data splitting was designed to randomly select 75% of the data within each of the indicated collections for fitting and 25% of the data within each of the indicated collections for testing. This design was chosen for consistency and comparability across the rows in Supplementary Fig. S2. When distinct DNA amounts were used for fitting and testing, the fitting and testing subsets were disjoint by nature. When common DNA amount(s) were used, 75% of the common data was (randomly) selected for fitting, while the rest was used for testing.

*Additional Figures*

(see next page)

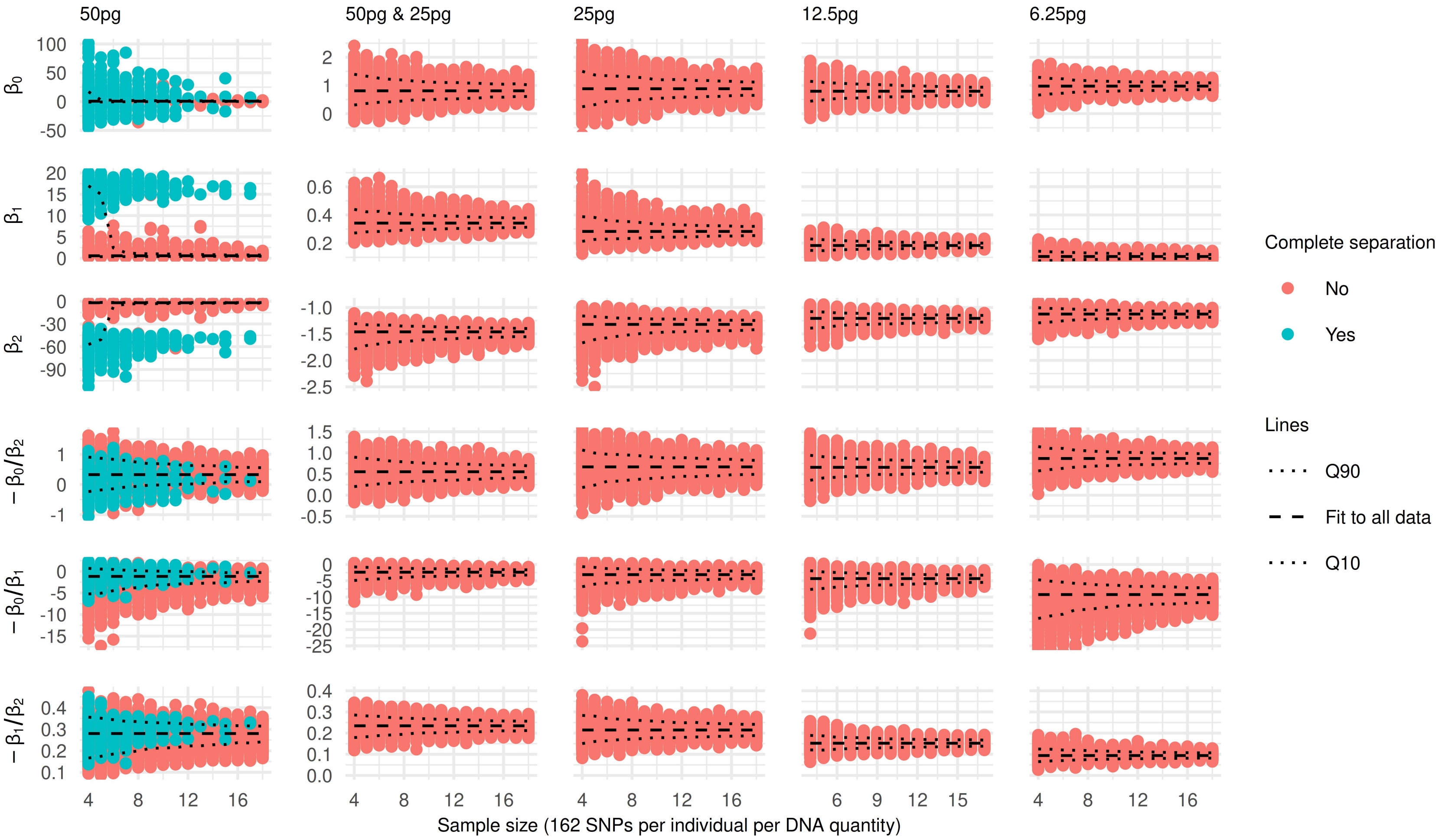


Supplementary Fig. S1: Parameter estimates from bootstrap analyses of the SMLR model across varying sample sizes.

The model was fitted with an intercept to square-root transformed allele signals from bootstrap samples drawn from the examinations of the DNA quantities indicated at the top of each plot column. The sample size refers to the number of sampled SNP profiles per indicated DNA quantity. Each SNP profile consists of 162 SNPs (including potential missing values). For each sample size, thousand bootstrap samples were drawn (with replacement), and for each of these, the SMLR model was fitted to obtain the parameter estimates indicated by red and blue dots. Blue dots indicate samples with complete separation (zero wrong calls), while red dots indicate samples without separation. Dashed lines represent the parameter estimates from fitting the model to all of the indicated data, and dotted lines show the 10th and 90th percentiles of the bootstrap estimates.

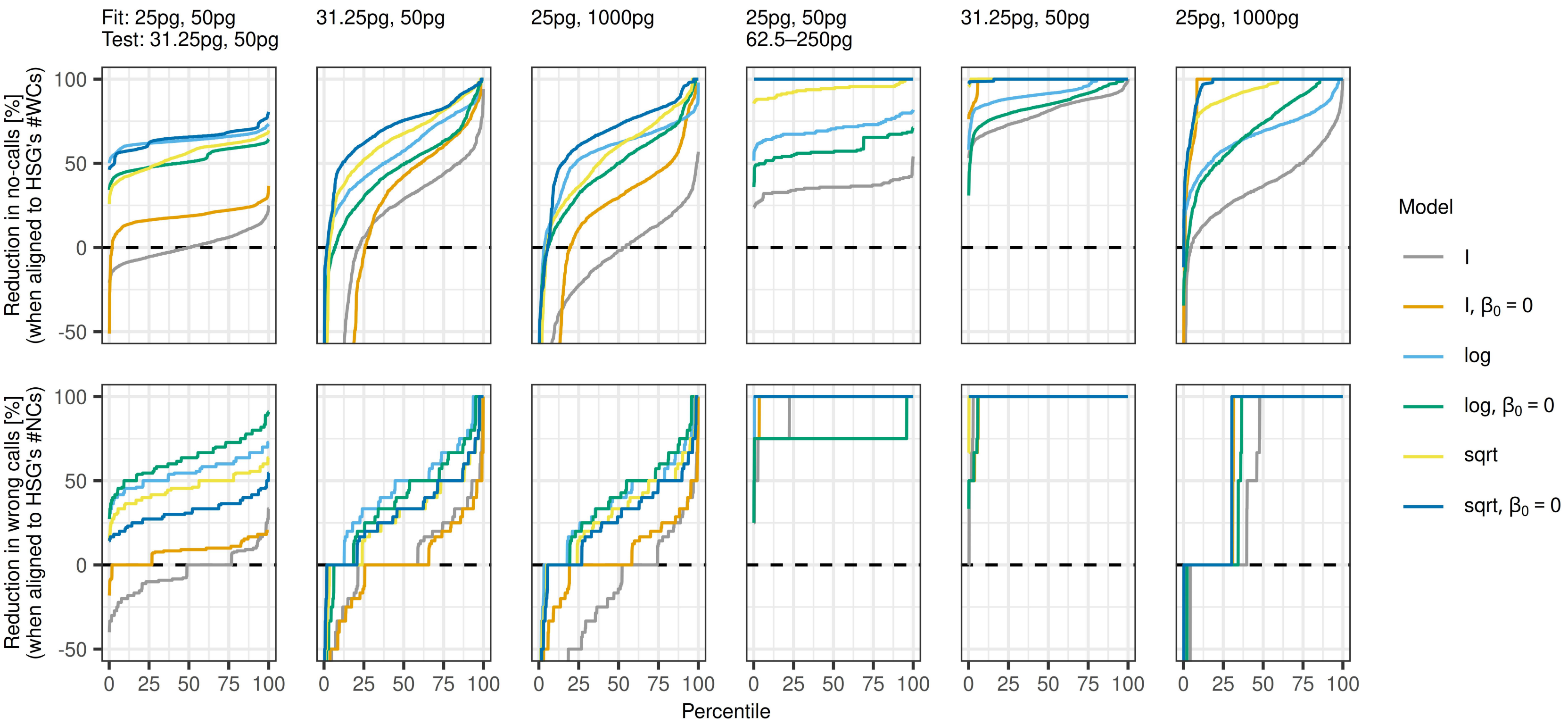


Supplementary Fig. S2: Performance comparisons of six variants of the SMLR model across different DNA amounts.

The models were evaluated using the metrics at (4) and (5) on each of 1,000 random splits of the data, each split dividing into 75% fitting data and 25% test data, with DNA amounts specified in the top. The first row of plots shows the SMLR models' reductions in no-calls (NCs) relative to the HID SNP Genotyper Plugin (HSG) when the SMLR models' no-call zones (in each data split) were set to the width where they made the same number of wrong calls (WCs) as the HSG or fewer. Similarly, the second row of plots shows the SMLR models' relative reductions in WCs when the no-call zones were set to the width where they made the same number of NCs as the HSG or fewer. In all subplots, the reductions are ordered and labelled according to their percentile.

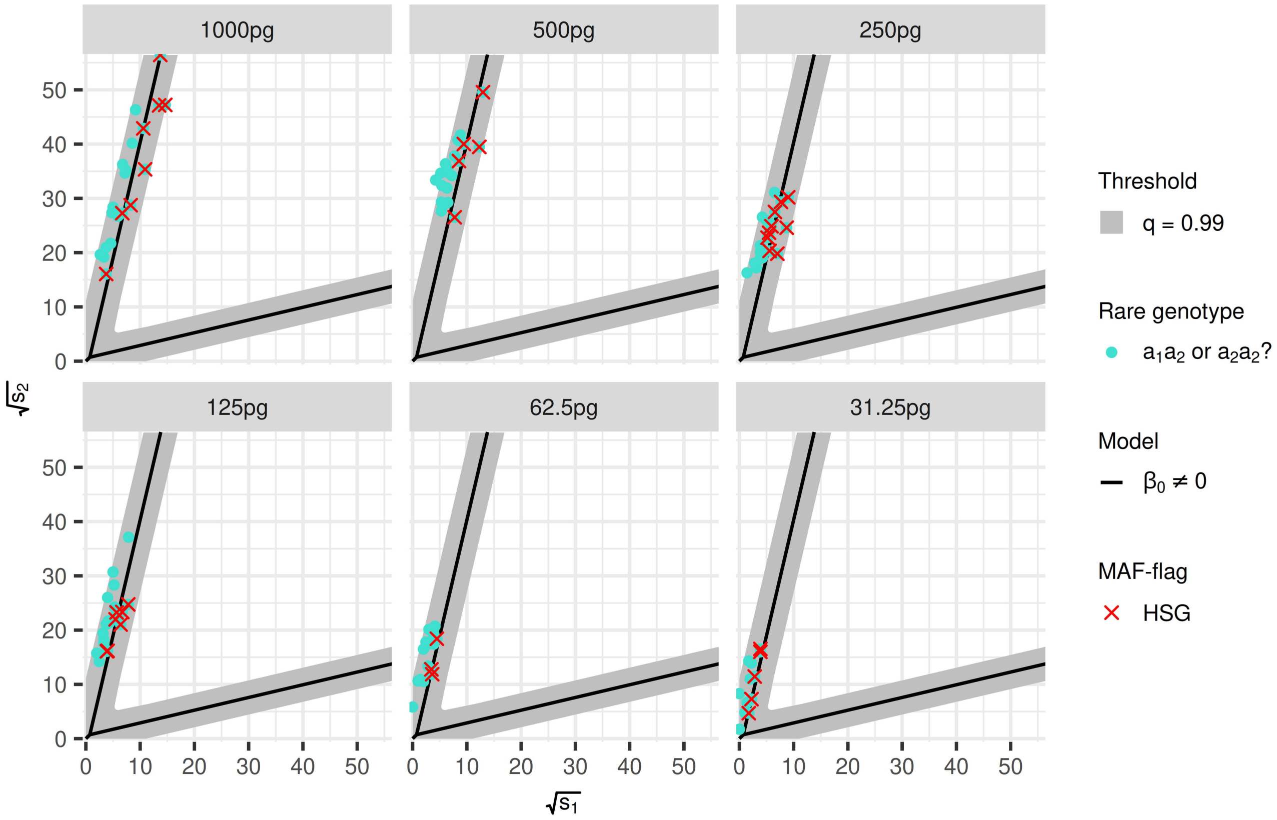


Supplementary Fig. S3: The SMLR framework used for quality check of the allele balance or spotting of rare alleles. The scatterplots show the major allele signal versus the adenine signal of the SNP locus rs7722456 for the DNA quantities examined in the first dilution series. The SMLR model (solid lines) was fitted with an intercept to square-root transformed allele signals from the examinations of 25 pg and 50 pg DNA from the second dilution series. Observations flagged by the HID SNP Genotyper Plugin for allelic imbalance concerns (MAF-flag) are marked with red crosses. The grey areas show where the posterior genotype probabilities of the SMLR model fall below a threshold value of $q = 0.99$, highlighting potential quality issues with the enclosed points.
